# The channel capacity of multilevel linguistic features constrains speech comprehension

**DOI:** 10.1101/2021.12.08.471750

**Authors:** Jérémy Giroud, Jacques Pesnot Lerousseau, François Pellegrino, Benjamin Morillon

## Abstract

Humans are expert at processing speech but how this feat is accomplished remains a major question in cognitive neuroscience. Capitalizing on the concept of channel capacity, we developed a unified measurement framework to investigate the respective influence of seven acoustic and linguistic features on speech comprehension, encompassing acoustic, sub-lexical, lexical and supra-lexical levels of description. We show that comprehension is independently impacted by all these features, but at varying degrees and with a clear dominance of the syllabic rate. Comparing comprehension of French words and sentences further reveals that when supra-lexical contextual information is present, the impact of all other features is dramatically reduced. Finally, we estimated the channel capacity associated with each linguistic feature and compared them with their generic distribution in natural speech. Our data point towards supra-lexical contextual information as the feature limiting the flow of natural speech. Overall, this study reveals how multilevel linguistic features constrain speech comprehension.

## Introduction

Humans are remarkably successful at quickly and effortlessly extracting meaning from spoken language. The classical method to study this ability and identify its processing steps is to reveal the constraints that limit speech comprehension. For example, the fact that speech comprehension drops when more than ∼12 syllables per second are presented has been interpreted as evidence that at least one processing step concerns syllables extraction (Ghitza, 2013; Giraud & Poeppel, 2012; Versfeld & Dreschler, 2002). As language processing involves distinct representational and temporal scales, it is usually decomposed into co-existing levels of information, estimated with distinct linguistic features, from acoustic to supra-lexical (Christiansen & Chater, 2016; Hickok & Poeppel, 2007; Rosen, 1992).

Recently, neuroimaging studies have started to incorporate simultaneously acoustic and linguistic features to model brain activity (e.g., Di Liberto et al., 2015; Cross et al., 2016). However, most speech comprehension studies, i.e. studies that include behavioral measures of language comprehension, only investigate a single linguistic feature and, as a consequence, a complete picture of which processes underlie speech comprehension is still lacking. This is because there exists no common theoretical framework and no unique experimental paradigm to compare multiple linguistic features at the same time. Among the existing experimental paradigms, artificially increasing the speaking rate to generate adverse and challenging comprehension situations is a common approach. However, when speech is artificially time-compressed (Dupoux & Green, 1997; Foulke & Sticht, 1969; Garvey, 1953), all linguistic features are impacted by the modification, making it impossible to disentangle their unique impact on behavioral performance. It thus remains unknown whether the syllabic rate actually constrains comprehension, whether it is the phonemic rate or any other rate, or whether bottlenecks are present at different levels of processing.

To solve this problem, we propose to rely on a concept inherited from information theory (Shannon, 1948), channel capacity, and to carefully orthogonalize multiple linguistic features to reveal their unique contribution to speech comprehension. The processing of each linguistic feature can be modeled as a transfer of information through a dedicated channel. Channel capacity is defined as the maximum rate at which information can be transmitted. Thanks to this approach, we identified and compared in a unique paradigm the potential impact of acoustic, sub-lexical, lexical and supra-lexical linguistic features on speech comprehension.

First, speech is an acoustic signal characterized by a prominent peak in its envelope modulation spectrum, around 4-5 Hz, a feature shared across languages (Ding et al., 2017; Varnet, Ortiz-Barajas, Erra, Gervain, & Lorenzi, 2017). This *acoustic modulation rate* approximates the *syllabic rate* of the speech stream (Poeppel & Assaneo, 2020), which happens at around 2.5 – 8 syllables per second in natural settings (Coupé, Oh, Dediu, & Pellegrino, 2019; Kendall, 2013; Pellegrino, Coupé, & Marsico, 2011). The acoustic modulation rate can serve as an acoustic guide for parsing syllables (Mermelstein, 1975). In addition to these, comprehension depends on the linguistic coding of phonemic details, necessitating parsing speech at the *phonemic rate* (Ghitza, 2011; Giraud & Poeppel, 2012; Hyafil, Fontolan, Kabdebon, Gutkin, & Giraud, 2015; Peelle & Davis, 2012; Poeppel, 2003; Stevens, 2002). We thus estimated three speech rates, the raw *acoustic modulation rate*, and the linguistically-motivated *syllabic and phonemic rates*.

Second, syllabic and phonemic sub-lexical units carry linguistic information. A description of speech in terms of linguistic information rates rather than speech rates could be more appropriate to understand how language is processed (Coupé et al., 2019; Pellegrino et al., 2011; Reed & Durlach, 1998). Moreover, the information rate (in bits/s), rather than an absolute informational value (in bits), is a more relevant dimensional space (Coupé et al., 2019), in accordance with the fact that neurocognitive ressources are best characterized as temporal bottlenecks (Hasson, Yang, Vallines, Heeger, & Rubin, 2008; Honey et al., 2012; Lerner, Honey, Katkov, & Hasson, 2014; Lerner, Honey, Silbert, & Hasson, 2011; Vagharchakian, Dehaene-Lambertz, Pallier, & Dehaene, 2012). Hence, we estimated *syllabic and phonemic informational rates*.

Third, at the lexical and supra-lexical levels, probabilistic constraints regulate language processing. It has been suggested that speech processing depends on predictive computations to guide the interpretation of incoming information. Predictions of upcoming individual words depend on both prior knowledge and contextual information (Brodbeck, Hong, & Simon, 2018; Donhauser & Baillet, 2020; Gagnepain, Henson, & Davis, 2012; Gwilliams, Linzen, Poeppel, & Marantz, 2018; Kutas, DeLong, & Smith, 2011; Pickering & Garrod, 2007; Sohoglu, Peelle, Carlyon, & Davis, 2012). Lexical (or word) frequency, the probabilistic knowledge about word occurrences, has a strong impact on lexical access time (Brysbaert, Lange, & Wijnendaele, 2000; Ferreira, Henderson, Anes, Weeks, & McFarlane, 1996). Hence, we estimated the context-independant or *static lexical surprise rate*, i.e., the amount of unexpectedness of word occurrences per second (see Methods). Additionally, recent models based on deep neural networks exploit contextual lexical information to predict brain activity during natural speech processing (Caucheteux, Gramfort, & King, 2021; Goldstein et al., 2020; Heilbron, Armeni, Schoffelen, Hagoort, & de Lange, 2020; Schrimpf et al., 2020). We used CamemBERT (Martin et al., 2020), a transformer model trained for the French language, to estimate the *contextual lexical surprise rate*, i.e., the lexical surprise rate predicted by the context provided by each sentence.

To reveal the efficiency of the speech comprehension system and estimate its capacity and limitations with unprecedented levels of granularity, we developed and combined three innovative experimental approaches: 1) First, we developed the *compressed speech gating paradigm*, a behavioral approach allowing an efficient estimation of the relation between time-compression and comprehension performance. For each stimulus a comprehension point could be determined, corresponding to the compression rate at which comprehension emerges. 2) Second, speech is in essence a temporal signal, and previous work has shown the relevance of considering linguistic features as a number of units communicated per unit of time (i.e., in rate, or bit/s; (Coupé et al., 2019; Pellegrino et al., 2011; Reed & Durlach, 1998). Each linguistic feature was thus expressed in a number of units per second. With such an approach, and utilizing the comprehension point as the maximum rate at which information is transmitted, the channel capacity associated with each linguistic feature can be estimated. Moreover, features can also be compared directly between one another and ranked according to the magnitude of their respective influence. 3) Third, to simultaneously estimate the impact of multiple linguistic features on comprehension capacities, we developed an original stimulus selection and orthogonalization procedure. We generated two speech corpora derived from large databases of natural stimuli and characterized them at the previously described seven linguistic levels, ranging from acoustic to supra-lexical. Thanks to a careful selection, all these features were orthogonalized across stimuli, enabling a fine-grained characterization of their respective influence on speech comprehension. The combination of these three methodological advances provides optimal conditions to investigate the linguistic features governing speech processing ability and limits.

Results from three behavioral experiments converge to show that multilevel linguistic features independently constrain speech comprehension, with the syllabic rate having the strongest impact. When supra-lexical contextual information is provided to participants, the impact of all other features is dramatically reduced. Estimating the channel capacity associated with each feature, we show in particular that comprehension drops when phonemic or syllabic rates are respectively above ∼40 Hz or ∼15 Hz. Finally, comparing these estimated channel capacities with the generic distribution of the linguistic features in natural speech, we find that at original speed contextual lexical information is already close to its channel capacity, which suggests that it is the main cognitive feature limiting the flow of natural speech.

## Results

### Compressed speech gating paradigm

We collected behavioral data from three independent experiments in which participants were required to understand successive time-compressed versions of either spoken monosyllabic words or sentences, respectively in Experiment 1, 2 and 3 (Fig. 1).21, 21 and 20 participants (age range: 20–43 years; 57% of females) were respectively recruited for experiments 1, 2 and 3. At each trial, the same spoken utterance was presented at decreasing compression rates ranging from unintelligible, to challenging, to intelligible. Using regression analyses, we modeled the individual comprehension performance fluctuation at the single trial level, as a function of a mixture of features encompassing the entire linguistic hierarchy from acoustic to supra-lexical levels of description. Linguistic features were chosen based on a large body of literature identifying them as influential constraints on speech comprehension (see Introduction). Our corpus selection procedure guaranteed that feature distributions selected in the final experimental material were representative of generic stimuli statistics as derived from large databases (Fig. Supp. 1a and 1d). In experiment 1, the limitations in terms of existing monosyllabic words prevented us from reaching a stimulus set in which the syllabic information rate was representative of the original database. Specifically, both the mean and variance of the distribution across stimuli differed between the original and selected stimulus sets (Fig. Sup. 1b and 1e). We thus excluded this feature from the data analyses of experiment 1. We also ensured that within the subset of selected stimuli, correlations between linguistic features were low (all r < 0.12; Fig. Supp. 1c and 1f), thanks to an orthogonalization procedure. This is a crucial condition to be able to determine their respective impact on speech comprehension performance. Finally, by investigating each feature in a similar measurement framework we were able to directly compare their respective impact on speech comprehension.

**Figure 1.**
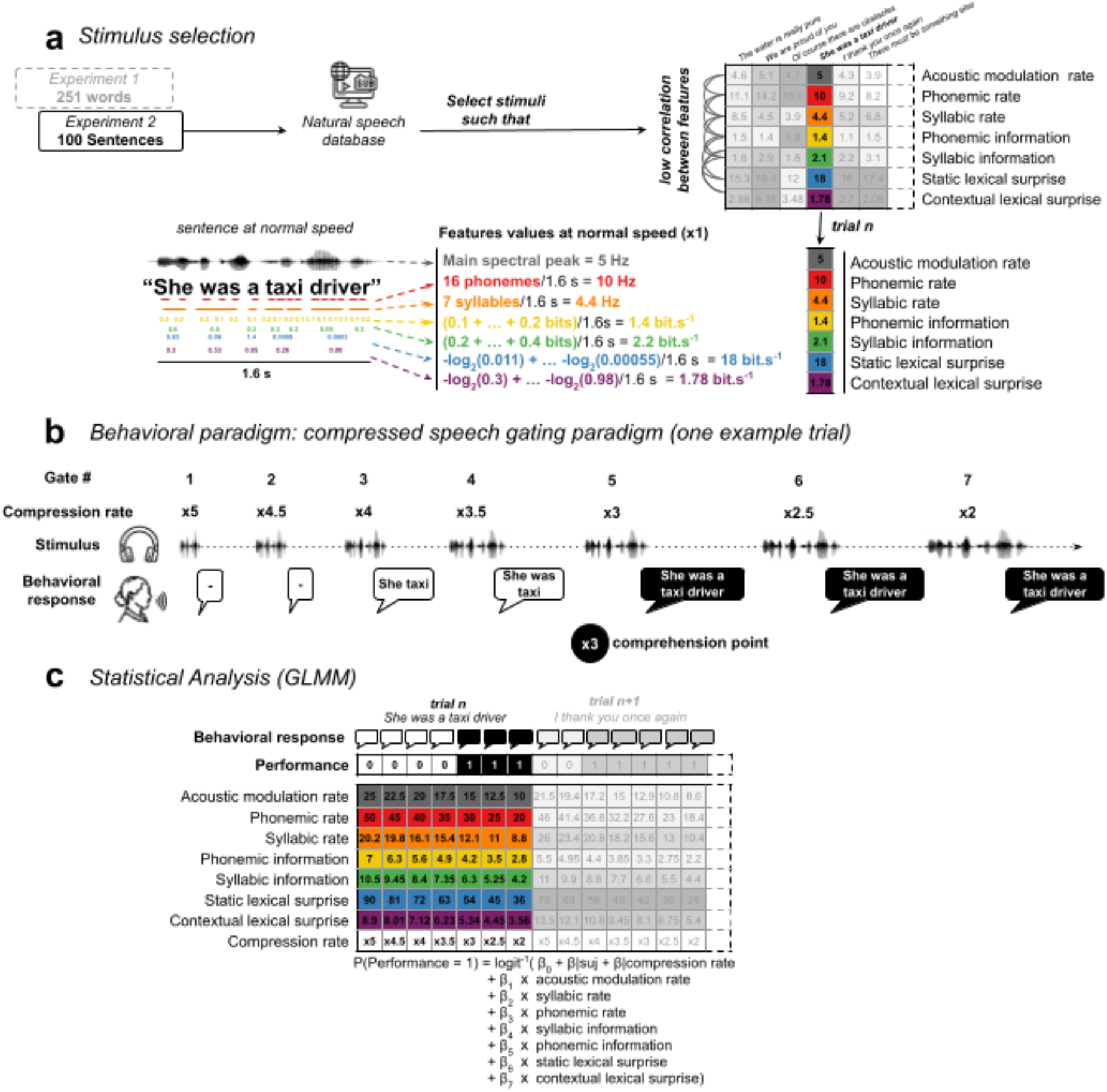
Experimental design and analysis pipeline. **a)** Stimulus selection procedure. 251 words and 100 sentences were used in experiments 1 and 2, respectively. Word stimuli were retrieved from the French Lexique database and sentence stimuli from the Web Inventory of Transcribed and Translated Talks database. Seven linguistic features were computed for each stimulus, illustrated here for an example sentence (sentences in experiment 2 were 7-words long). Features corresponded to the acoustic modulation rate (in Hz), syllabic rate (in Hz), phonemic rate (in Hz), syllabic information rate (in bit/s), phonemic information rate (in bit/s), static lexical surprise (in bit/s) and contextual lexical surprise (in bit/s). The selection procedure ensured that low correlations (all r < 0.15) across stimuli were present between features in the selected stimulus sets (see Methods). **b)** Behavioral paradigm. A modified gating paradigm was used for both experiments. In each trial, participants were presented with time-compressed versions of the original audio stimulus, from the most to the least compressed version, and were asked to report what they heard after each audio presentation. Behavioral responses were classified into incorrect and correct responses (incorrect: white bubbles; correct: black bubbles). At each trial, a “comprehension point” (black circle) was determined. It corresponds to the compression rate at which comprehension emerged, estimated across gates with a logistic regression model (see Methods). **c)** Behavioral responses were entered into a generalized linear mixed models (GLMM) to assess the respective contribution of each feature on comprehension performance. The equation includes participants and compression rates as random effects and linguistic features as fixed effects. Entering compression rates as random effects ensured that correlations between stimuli across compression rates were controlled for in the model.

### Compressed speech impairs speech comprehension

Across the different compression rates, comprehension shifted from not understood (mean performance accuracy of 0.03 % and 0.1 % for experiments 1 and 2, respectively) to perfectly understood (96.3 % and 99 %), with a characteristic sigmoid function, indicating that the range of compression rates selected was well suited to investigate speech comprehension at its limits (Fig. 2). A mean performance accuracy of 50 % was observed for a compression rate of 3.5 in both experiments. At a compression rate of 5 or above, comprehension was essentially residual (< 10 %).

**Figure 2.**
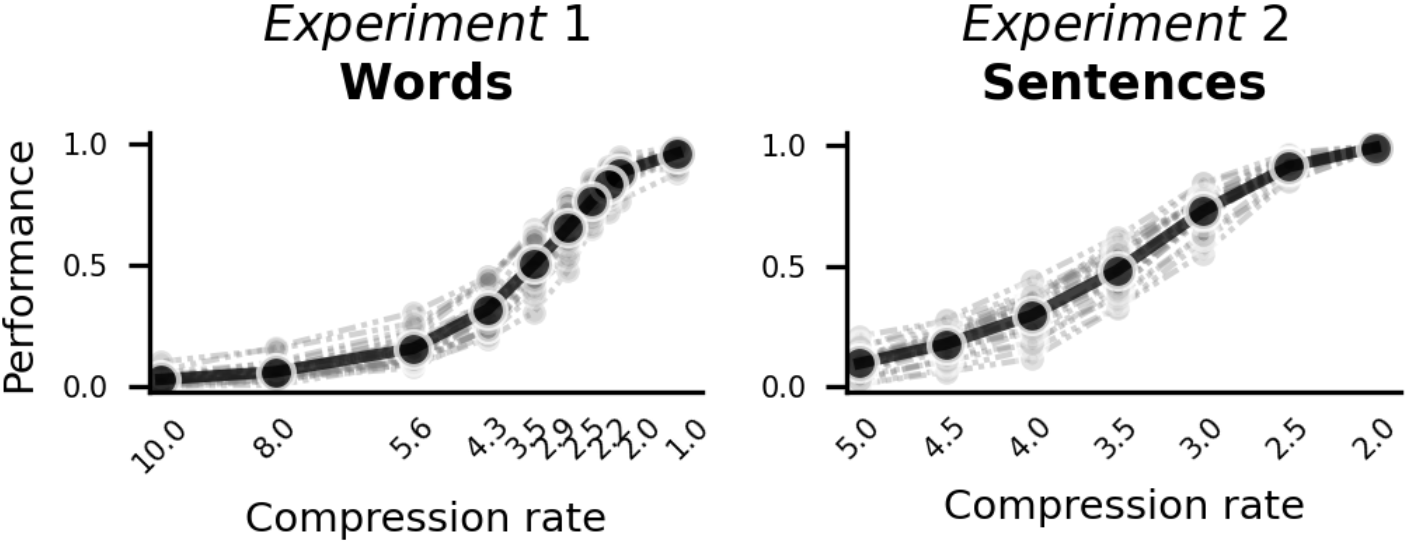
Comprehension performance as a function of compression rate. Performance is expressed in proportion of correct responses. Thin dashed grey lines depict individual performance. Thick black lines indicate average performance. In experiment 1, participants were presented with the same audio stimuli (words) at ten different compression rates. In experiment 2, participants were presented with the same audio stimuli (sentences) at seven different compression rates.

### Multifactorial linguistic constraints concurrently limit speech comprehension

We used generalized linear mixed-effects models (GLMMs) to evaluate the extent to which multiple linguistic features were predictive of behavioral performance (word or sentence comprehension). The GLMM approach enables a fine-grained characterization of the independent contributions of the different features (see Methods).

In experiment 1, a GLMM with a logit link function was conducted to model spoken word comprehension. The model included participants and compression rates as random effects and five linguistic features, acoustic modulation rate, the phonemic and syllabic rates, phonemic information rate and static lexical surprise, as fixed effects (Fig. 3, left panel; table 1; see Methods). The stimuli consisting of isolated words, no contextual lexical surprise was defined. The full model accounted for 74 % of the variance of the data. The model revealed a significant effect of the acoustic modulation rate (β = -0.7 ± 0.06, p < 0.001), the phonemic rate (β = -0.25 ± 0.07, p = 0.001) and the syllabic rate (β = -1.07 ± 0.08, p < 0.001), indicating that they independently and additively impact comprehension. The model’s coefficients read as follows:β = -1.07 means that the odds of giving a correct response are multiplied by exp(−1.07) ≈ are divided by 3 for an increase of one standard deviation in syllabic rate, demonstrating the adverse impact of increased syllabic rate on speech comprehension. Phonemic information rate did not significantly contribute to the model (β = -0.03 ± 0.03, p = 0.258). Finally, the static lexical surprise was significantly associated with listeners’ speech comprehension (β = -0.91 ± 0.07, p < 0.001), indicating that words’ unexpectedness worsens participants’ comprehension.

**Figure 3.**
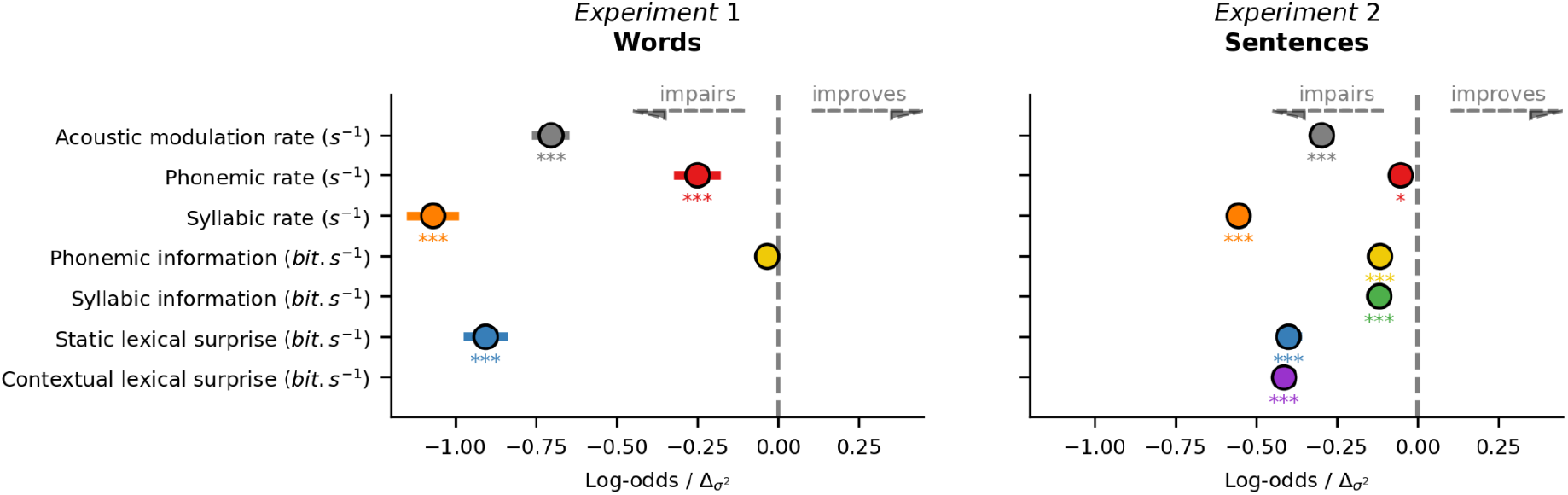
GLMM results. Log-odds ratios of the linguistic features included in the GLMM models in experiments 1 and 2. Coefficients were standardized and read as follows: in experiment 1, the odds of giving a correct response are multiplied by exp(−1.07) ≈ 0.33 ≈ are divided by 3 for an increase of one standard deviation in syllabic rate (orange dot in experiment 1). In other words, an increase of one standard deviation in syllabic rate divides the odds of understanding the word by 1/exp(−1.07) ≈ 3. Negative log-odds ratios indicate a negative effect on performance. In both models, linguistic features were entered as fixed effects. Participants and compression rates were entered as random effects. *p < 0.05; ***p < 0.001. Error bars indicate standard error of the mean across participants.

**Table 1.** Results from the Generalized (binomial) Linear Mixed Models for experiments 1 and 2 with comprehension performance as dependent variable. Acoustic modulation rate, phonemic rate, syllabic rate, phonemic information rate, syllabic information rate, static lexical surprise and contextual lexical surprise as fixed effects in experiment 2 model. In experiment 1 model, syllabic information rate and contextual lexical surprise are not included. All fixed effects were z-transformed to obtain comparable estimates. Random intercepts are also included for each participant. 10 and 7 compression rates are included as random variables in experiment 1 and experiment 2 respectively. 21 participants took part in experiment 1 and 21 participants took part in experiment 2. The models were run on 52710 and 14700 individual responses in experiment 1 and 2 respectively. Statistical significance of predictors was assessed using likelihood ratio tests (p).

|  | Experiment 1 (words) |  |  |  |  | Experiment 2 (sentences) |  |  |  |  |
| --- | --- | --- | --- | --- | --- | --- | --- | --- | --- | --- |
| Fixed effects |  |  |  |  |  |  |  |  |  |  |
|  | Log-odds | SE | CI (95%) |  | p | Log-odds | SE | CI (95%) |  | p |
| Intercept | -0.02 | 0.32 | -0.64 | 0.60 | 0.956 | 0.43 | 0.31 | -0.17 | 1.04 | 0.161 |
| Acoustic modulation rate | -0.70 | 0.06 | -0.82 | 0.59 | **<0.001** | -0.30 | 0.03 | -0.35 | -0.24 | **<0.001** |
| Phonemic rate | -0.25 | 0.07 | -0.39 | -0.11 | **0.001** | -0.05 | 0.02 | -0.10 | -0.01 | **0.022** |
| Syllabic rate | -1.07 | 0.08 | -1.23 | -0.91 | **<0.001** | -0.56 | 0.02 | -0.60 | -0.51 | **<0.001** |
| Phonemic information rate | -0.03 | 0.03 | -0.10 | 0.03 | 0.258 | -0.12 | 0.02 | -0.16 | -0.07 | **<0.001** |
| Syllabic information rate |  |  |  |  |  | -0.12 | 0.02 | -0.16 | -0.08 | **<0.001** |
| Static lexical surprise | -0.91 | 0.07 | -1.04 | -0.77 | **<0.001** | -0.40 | 0.04 | -0.48 | -0.32 | **<0.001** |
| Contextual lexical surprise |  |  |  |  |  | -0.41 | 0.01 | -0.44 | -0.39 | **<0.001** |
| Random effects |  |  |  |  |  |  |  |  |  |  |
| $\sigma^2$ | | | 3.29 | | | | | 3.29 | | |
| $T_{00}$ | | | 0.14 participant<br>0.93 compression rate | | | | | 0.16 participant<br>0.62 compression rate | | |
| ICC |  |  | 0.25 |  |  |  |  | 0.19 |  |  |
| Number of observations |  |  |  |  |  |  |  |  |  |  |
| N |  |  | 10 compression rate<br>21 participant |  |  |  |  | 7 compression rate<br>21 participant |  |  |
| Observations |  |  | 52710 |  |  |  |  | 14700 |  |  |
| Marginal R<sup>2</sup> / Conditional R<sup>2</sup> |  |  | 0.659 / 0.743 |  |  |  |  | 0.427 / 0.536 |  |  |

Holm-corrected post-hoc comparisons were performed to identify differences among selected features in modulating spoken word comprehension. Features were ordered from the most to the least influential, and compared between neighbours. The analysis revealed no significant difference between the two most influential features, syllabic rate and static lexical surprise (β = - 0.16, z = -1.58, p = 0.12). In contrast, all other pairwise comparisons were significantly different (all p < 0.05).

In experiment 2, a GLMM with a logit link function was also used to model spoken sentences comprehension. The model included seven linguistic features as fixed effects (Fig. 3, right panel; table 1; see Methods). All linguistic features significantly contributed to the model and together explain 54 % of the variance of the data (Fig. 3, right panel; table 1). Similar to experiment 1, post-hoc comparisons were conducted to assess differences between the relative influence of each linguistic feature on sentence comprehension. The analysis showed that the syllabic rate has the largest impact on performance, with significantly more influence than contextual lexical surprise (β = -0.14, z = -5.22, p < 0.001). Conversely, the contrast between contextual and static lexical surprise rate did not reach significance (β = -0.01, z = -0.34, p > 0.05). Whereas modulatory effect of the static lexical surprise and the acoustic modulation rate on comprehension was not significantly different (β = -0.10, z = -2.07, p > 0.05), this latter alter significantly more speech comprehension than syllabic information rate (β = -0.18, z = -4.87, p < 0.001). Finally, modulation of performance induced by syllabic information rate, phonemic information rate and phonemic rate do not significantly differ (all p > 0.41).

### Adding contextual information reduces the influence of the other linguistic features

Comparing experiments 1 and 2, we first observed a similar profile of response weights, with a larger impact of syllabic rate and static lexical surprise, a medium influence of the acoustic modulation rate, and lower weights for the other linguistic features (Fig. 3).

We assessed, for each linguistic feature, potential significant differences between experiments 1 and 2. This analysis (Fig. Supp. 3) reveals that the weights associated with the four features of interest - the acoustic modulation, phonemic and syllabic rates and the static lexical surprise- are significantly larger in Experiment 1 than in Experiment 2 (all p < 0.05 Holm-corrected for multiple comparison). This difference is associated with a reduction of (around or more than) 50% in experiment 2 compared to experiment 1. This hence suggests that adding contextual lexical information (the main difference between experiments 1 and 2) reduces the impact of all other features on comprehension.

Of note, a fifth feature investigated in this comparison -phonemic information- was associated with a non-significant weight in experiment 1, a significant but marginal weight in experiment 2, and these weights are not significantly different across experiments, which confirms the marginal impact of this linguistic feature on comprehension.

### Multilevel linguistic features consistently shift the comprehension point

Following the main GLMM analysis, we aimed at characterizing the relationship between the value of each linguistic feature at original speed (x1) – which reflects the intrinsic linguistic properties of the stimulus sets – and the comprehension point (i.e the compression rate at which participants’ comprehension reaches 75 % of accuracy, see Methods). This analysis ought to confirm the individual propensity of each linguistic feature to modulate the comprehension point (see Methods). In experiment 1, a linear mixed model analysis fully reproduced the results from the main GLMM analysis (Table 2), revealing a significant impact of all features but the phonemic information rate, on comprehension (all p < 0.05). In experiment 2, the linear mixed model revealed that, apart from phonemic information rate, all other features significantly delayed the comprehension point (all p < 0.05), also confirming the previous analysis. The putative effect size associated with phonemic information rate is probably negligible, even if significance has been limited by the number of observations taken into account in this alternative model (2100 vs. 14700 behavioral responses, see Methods). Overall, these new analyses confirm the robustness of the results previously obtained with the GLMM and directly show that the linguistic properties of the non-compressed stimuli predict the maximal compression rate at which comprehension can be maintained.

**Table 2.** Results from the binomial Linear Mixed Models for experiment 1 and 2 with comprehension point as dependent variable. Acoustic modulation rate, phonemic rate, syllabic rate, phonemic information rate, syllabic information rate, static lexical surprise and contextual lexical surprise were entered as fixed effects in experiment 2 model. In experiment 1 model, syllabic information rate and contextual lexical surprise are not included. All fixed effects were z-transformed to obtain comparable estimates. Random intercepts are also included for each participant. The models were run on 5250 and 2100 individual responses in experiment 1 and 2 respectively. Statistical significance of predictors was assessed using likelihood ratio tests (p).

|  | Experiment 1 (words) |  |  |  |  | Experiment 2 (sentences) |  |  |  |  |
| --- | --- | --- | --- | --- | --- | --- | --- | --- | --- | --- |
| Fixed effects |  |  |  |  |  |  |  |  |  |  |
|  | Log-odds | SE | CI (95%) |  | p | Log-odds | SE | CI (95%) |  | p |
| Intercept | 3.92 | 0.10 | 3.73 | 4.10 | <0.001 | 3.23 | 0.03 | 3.16 | 3.29 | <0.001 |
| Acoustic modulation rate | -0.19 | 0.03 | -0.25 | -0.13 | <0.001 | -0.03 | 0.01 | -0.06 | -0.01 | 0.002 |
| Phonemic rate | -0.06 | 0.03 | -0.12 | -0.01 | 0.027 | -0.03 | 0.01 | -0.05 | -0.00 | 0.019 |
| Syllabic rate | -0.21 | 0.03 | -0.26 | -0.15 | <0.001 | -0.05 | 0.01 | -0.08 | -0.03 | <0.001 |
| Phonemic information rate | -0.01 | 0.03 | -0.06 | 0.05 | 0.765 | 0.00 | 0.01 | -0.02 | 0.02 | 0.989 |
| Syllabic information rate |  |  |  |  |  | -0.03 | 0.01 | -0.06 | -0.01 | 0.002 |
| Static lexical surprise | -0.20 | 0.03 | -0.26 | -0.15 | <0.001 | -0.04 | 0.01 | -0.07 | -0.02 | <0.001 |
| Contextual lexical surprise |  |  |  |  |  | -0.20 | 0.01 | -0.22 | -0.18 | <0.001 |

**Table 2.** Results from the binomial Linear Mixed Models for experiment 1 and 2 with comprehension point as dependent variable.
| Random effects |  |  |
| --- | --- | --- |
| $\sigma^2$ | 4.16 | 0.25 |
| $\tau_{00}$ | 0.18 participant | 0.02 participant |
| ICC | 0.04 | 0.08 |
| Number of observations |  |  |
| N | 21 participant | 21 participant |
| Observations | 5250 | 2100 |
| Marginal $R^2$ / Conditional $R^2$ | 0.025 / 0.065 | 0.172 / 0.241 |

### The syllabic rate is the strongest determinant of speech comprehension

To more directly visualise the data from both experiments, a complementary approach was adopted. For each compression rate, performance was first binned as a function of the syllabic rate (see Methods), as this feature had the strongest impact on performance in the two experiments (Fig. Supp. 3 and Fig. Supp. 4a). This visualisation highlights the major influence of the syllabic rate on behavioral outcome independently of the compression rate, in both experiments. Second, data were also binned as a function of the other features, after having been stratified as a function of the syllabic rate (Fig. Supp. 3 and Fig. Supp. 4). This highlights their additional impact over the major influence of syllabic rate. This visualisation enables a better grasping of the relative influence of each linguistic feature on comprehension and confirmed graphically the genuine results obtained with the more fine-grained GLMM and LMM approaches.

### Stimulus repetition has no effect on comprehension performance

The compressed speech gating paradigm requires that the same speech stimulus be repeated immediately with a lower compression rate. Such a procedure could bias the comprehension point in favour of earlier comprehension, as participants might understand a little more with each repetition of the stimulus. Although this paradigm specificity is unlikely to have an impact on the main results (e.g. GLMM/LMM analyses, Fig. 3), it is possible that the comprehension point would occur later if the stimuli were not repeated immediately.

In order to address this concern, we ran a control experiment (experiment 3). We recruited a new pool of twenty participants online. They performed a shorter version of experiment 2. The participants were presented with the same stimuli than in experiment 2, but at only one compression rate (*3.5), the compression rate leading to approximately 50% of comprehension in experiment 2 (the inflexion point of the sigmoid curve of comprehension). Importantly, in experiment 2, this compression rate corresponded to the gate n°4, i.e., the fourth repetition of the same sentence in a row, while in the new experiment it corresponds to the first and unique presentation (gate n°1). It is hence appropriate to investigate the potential impact of stimulus repetition on comprehension. Like in experiment 2, participants were asked to repeat the sentence after each single presentation. Data were scored exactly as in experiment 2.

We assessed whether stimulus repetition was biasing the comprehension point and our estimation of the channel capacities associated with each linguistic feature. We performed an independent t-test to assess the difference of performance between experiments 2 and 3. The statistical procedure revealed no significant difference between the two samples (p> 0.05, t(39) = - 1.8; Fig. 4), which indicates that stimulus repetition does not facilitate comprehension compared to a unique presentation nor bias the comprehension point towards earlier understanding, and hence does not bias our estimation of the channel capacities associated to each linguistic features.

**Figure 4.**
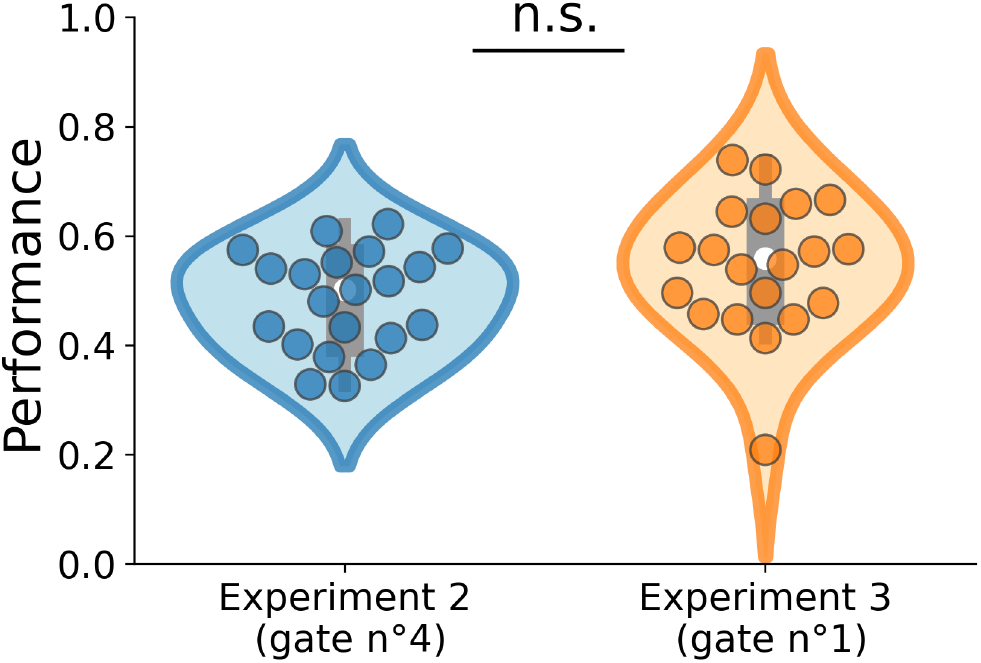
Mean individual comprehension performance in experiment 2 and 3 obtained at compression rate *3.5. In both experiments, the same sentence stimuli were presented at the same compression rate (*3.5). In experiment 2, it corresponded to the 4^th^ gate (4^th^ repetition) whereas in experiment 3, it was the first time that participants were presented with the stimuli (1^rst^ gate). An independent t-test reveals no significant difference in performance across experiments (p > 0.05, t(39)= -1.8). This result indicates that in our original experiments, repetition does not bias the comprehension points and hence that our estimation of the channel capacities associated to each linguistic feature is accurate.

To summarize, the repeated presentation paradigm (experiment 2) and the unique presentation paradigm (experiment 3) yield converging estimations in terms of linguistic feature importance and channel capacity estimation.

### Estimation of the channel capacity associated with each linguistic feature

Thanks to the compressed speech gating paradigm, we were able to derive for each feature the distribution of its values (in rate) at the comprehension point, which provided an estimation of its channel capacity (see Methods). This estimation corresponds to the value (in rate, or bit/s) at which comprehension consistently emerges. This threshold thus reflects a successful transmission of linguistic information but also determines the highest rate of information flow. As such, stimuli containing linguistic feature’s values above this threshold will exceed channel capacity leading to a drop in comprehension performance. Overall, we found that channel capacities associated with each linguistic feature investigated were on the same order of magnitude in both experiments (Fig. 5). Specifically, the estimated maximum acoustic modulation and syllabic rates were both centred around 10-15 Hz, while the phonemic rate’s channel capacity was centred around 35 Hz.

**Figure 5.**
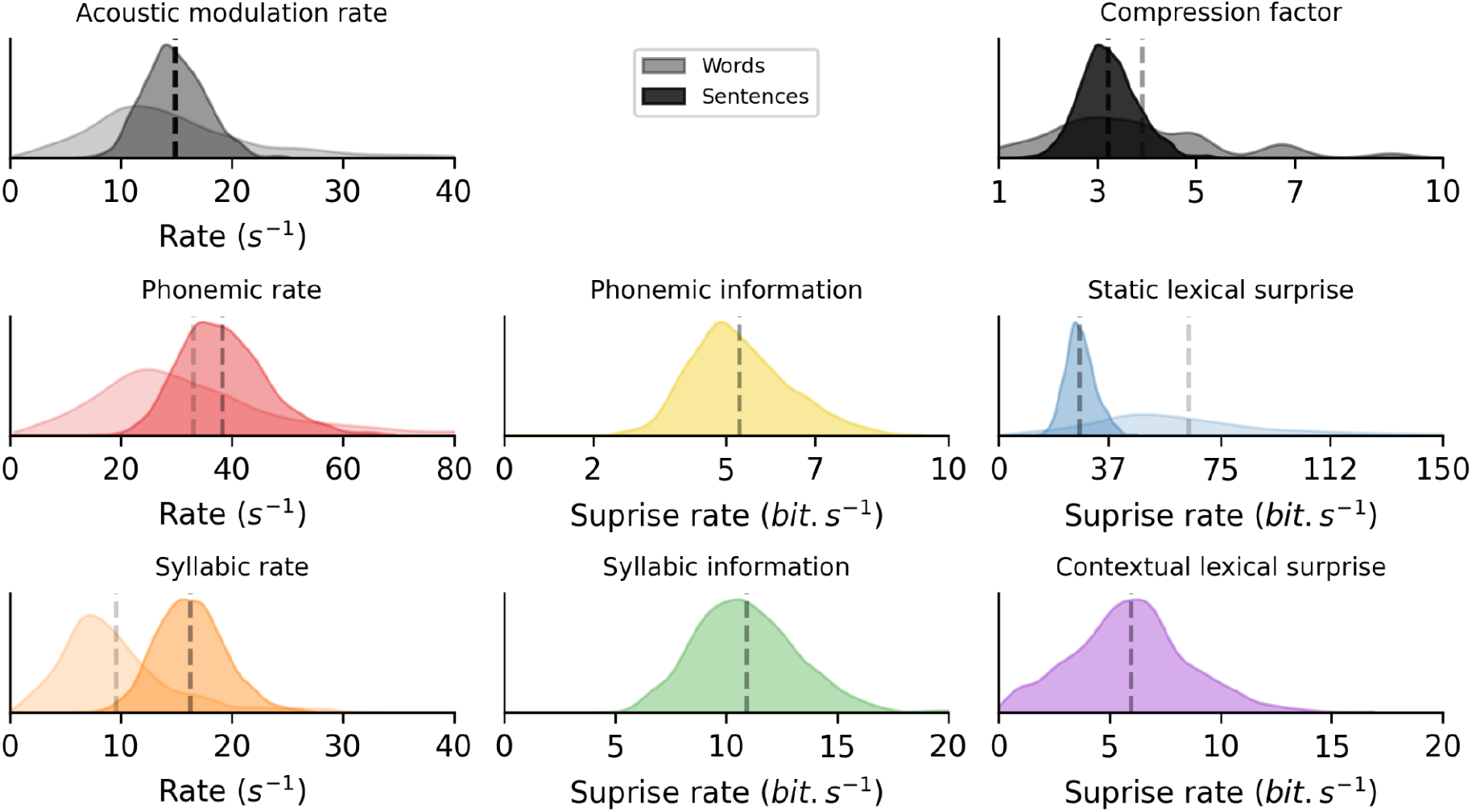
Channel capacity associated with each linguistic feature estimated in experiments 1 (words) and 2 (sentences). At each trial, the comprehension point – which corresponds to the compression rate at which comprehension emerged – was estimated (upper right panel, see Methods). As each feature significantly impacts comprehension (see Fig. 3), their maximal rate before they begin to negatively impact comprehension can be estimated. Values of each linguistic feature at comprehension points were extracted and aggregated across trials. The resulting distribution provides an estimate of the channel capacity associated with each linguistic feature. Data from experiment 1 (words) is depicted in lighter colors. For each linguistic feature, the channel capacity estimated in experiments 1 and 2 are of the same order of magnitude. Dashed vertical lines indicate the median of each distribution.

### Contextual information rate constrains the flow of natural speech

We finally estimated whether any linguistic feature was close to its channel capacity in the non-compressed stimulus sets. For each linguistic feature, we thus compared its value at the comprehension point (*i*.*e*. its channel capacity) and at original speed (*i*.*e*. its intrinsic statistics) and estimated a percentage of overlap across distributions.

In experiment 2, for each feature, the percentage of overlap between the two distributions was below 1 %, with the exception of the contextual lexical surprise, which was reaching a ∼18 % of overlap (a value significantly higher than the others; repeated-measures ANOVA: F (6,140) = 3482.3, p < 0.001; post-hoc paired t-tests: contextual lexical surprise vs. others: all p < 0.001 Tukey- corrected; all other comparisons: p > 0.9 Tukey-corrected; Fig. 6, upper right panel). This indicates that it is not unusual in natural speech to observe an amount of contextual lexical surprise close to its channel capacity, while natural speech operates much farther from the channel capacity of the other linguistic features. In experiment 1, the percentage of overlap was around 5% for all features (repeated-measures ANOVA: F (3,80) = 4.9, p = 0.003; post-hoc paired t-tests, all p > 0.001 Tukey- corrected; Fig. Supp. 5).

**Figure 6.**
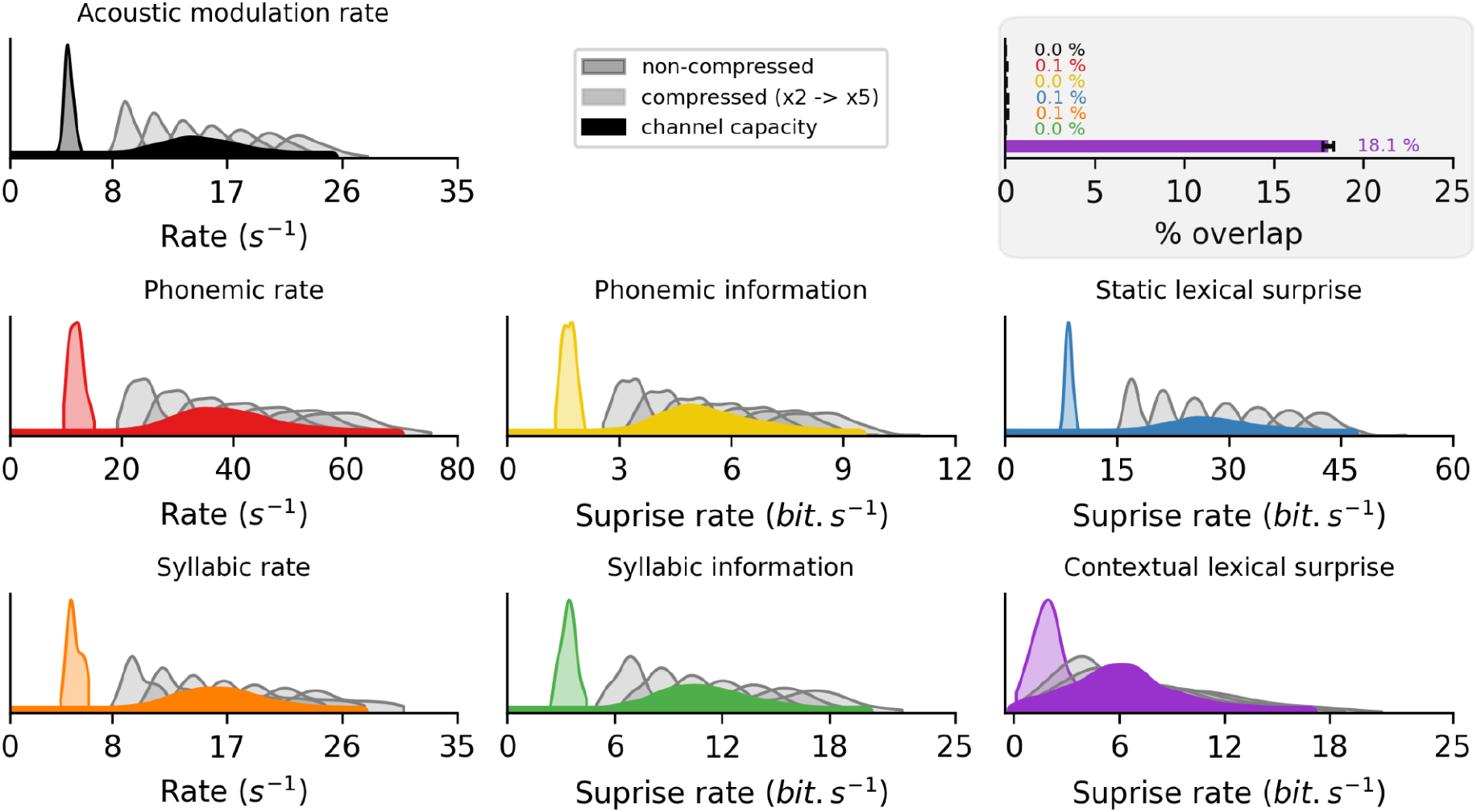
Experiment 2 (sentences). Overlap between the channel capacity associated with each linguistic feature and their generic distribution in the stimulus set. Distribution of the linguistic features in the selected stimulus set at original speed (non-compressed, lighter color) and at the different compression rates (in grey). Superimposed is their corresponding estimated channel capacity (see Fig. 5; darker color). **Upper right (grey panel):** Overlap ratio between the channel capacity associated with each linguistic feature and its generic distribution at original speed. Error bars indicate standard error of the mean across participants.

## Discussion

In this study, we investigated the extent to which multilevel linguistic features independently constrain speech comprehension. We expressed each linguistic feature in a number of units per second and derived their associated channel capacity thanks to an innovative experimental paradigm, the compressed speech gating paradigm. Guided by previous lines of research on speech comprehension (Coupé et al., 2019; Ghitza, 2014; Giraud & Poeppel, 2012; Schrimpf et al., 2020), we focused on features encompassing the entire linguistic hierarchy, from acoustic to supra-lexical levels of description, and investigated their individual effect on trial-by-trial performance fluctuations using generalized mixed linear model (GLMM) analyses. We report convergent results using two independent sets of stimuli (words and sentences) and participant sets. Moreover, we showed the robustness of the findings across two different experimental settings (in-lab and online) and complementary analyses (GLMM and LMM). Finally, we reproduce key findings from the literature and report plausible conclusions, compatible with current theoretical models and known biological evidence.

Previous work has focussed on characterizing prominent speech features relevant for comprehension. In particular, speech has been described as an inherently rhythmic phenomenon, in which linguistic information is pseudo-rhythmically transmitted in “packets” (Ghitza, 2014). The theta timescale (4-8 Hz), associated with the main acoustic modulation and the syllabic rates, has been highlighted for its main contribution to speech comprehension (Ahissar et al., 2001; Poeppel & Assaneo, 2020). Moreover, speech-specific temporal organisation is thought to be reflective of an evolutionary attempt to maximize information transfer given cognitive and neural constraints (Christiansen & Chater, 2016). Accordingly, recent experimental evidence suggests that despite multiple differences, languages are highly similar in terms of average rate of transmission of information (Coupé et al., 2019). Our work is a critical extension of these previous lines of research, by directly comparing multiple relevant features and timescales for speech comprehension into a common measurement framework.

We first behaviorally confirmed human impressive ability to cope with highly speeded speech but also showed a collapse of language comprehension when spoken stimuli presentation rate exceeds a given threshold, i.e. beyond a compression factor of 3 (Dupoux & Green, 1997; Foulke & Sticht, 1969; Ghitza, 2014; Nourski et al., 2009). We show that this phenomenon can be explained as the result of a linear combination of multiple processing bottlenecks along the linguistic hierarchy. Corroborating previous findings, we show that the syllabic rate is the strongest determinant of speech comprehension.

Theoretical models propose that speech is sampled in parallel at two timescales, corresponding to the syllabic and phonemic rates (Giraud & Poeppel, 2012). To date, experimental evidence only established that specific brain rhythms in the auditory cortex track the acoustic dynamics during speech perception (Gross et al., 2013; Luo & Poeppel, 2007; Peelle, Gross, & Davis, 2013). Here we directly extended these results at the perceptual level by testing the impacts of the acoustic modulation, syllabic and phonemic rates on comprehension with a tightly orthogonalized setup. Our data reveal that these three features independently constrain speech comprehension. In particular, we found that channel capacities associated with acoustic modulation and syllabic rates were at around 15 Hz while the channel capacity associated with the phonemic rate was at around 35 Hz. These values parallel theoretical considerations and neurophysiological observations (Giraud & Poeppel, 2012; Giroud et al., 2020) and provide a behavioral validation that phonemic sampling occurs at such a rate (see also (Marchesotti et al., 2020). While the acoustic modulation and syllabic rates are often reduced to one another, they are dissociable (see also (Schmidt et al., 2021), are associated with different processing bottlenecks, but both unfold at around 5 Hz in natural speech and have a channel capacity of around 15 Hz. This result strongly suggests that both low-level acoustic and language-specific rhythmic processes contribute to speech comprehension. The channel capacities estimated for higher-order linguistic features cannot be compared with anything currently known in the literature. These results provide directly testable hypotheses for future human neurophysiology experiments.

French has been described as a syllable-timed language or, as Laver rightly nuanced, syllable-based (Laver, 1994). However, recent corpus-based studies revealed a high variability (Arvaniti, 2009; Barry, Andreeva, & Koreman, 2009; Jadoul, Ravignani, Thompson, Filippi, & de Boer, 2016; Wiget et al., 2010) and as a result, the idea of a strict categorical distinction between stress-timed and syllable-timed languages has now been discredited (Payne, 2021); see also (Rathcke & Smith, 2015). Critically, experimental works in various languages have highlighted the fundamental role of the syllable in speech perception, independently of the ‘category’ (syllable- or stress-based) of the investigated language, the syllabic rate being: (1) similar across languages (Coupé et al., 2019; Ding et al., 2017; Varnet et al., 2017); (2) at the foundation of speech segmentation (Poeppel & Assaneo, 2020; Strauß & Schwartz, 2017); and (3) a strong determinant of speech comprehension across languages (Ghitza & Greenberg, 2009; Ghitza, 2012; Versfeld & Dreschler, 2002). Overall, these findings support the view that our results can be generalized to “non syllable-timed” languages.

Additionally, by developing a normative measurement framework, we bridged speech perception studies with the domains of psycholinguistics, computational linguistics and natural language processing. First, our data reveal a mild adversarial effect of information rate at the phonemic and syllabic scales on speech comprehension. Whether these effects are similar across languages remains an open question. However, previous experimental evidence supports the view that the channel capacities that we estimated would reflect the general human cognitive architecture or the ecological language niche (Coupé et al., 2019; Pellegrino et al., 2011). Second, we show that the respective impact on comprehension of the syllabic rate, the static lexical surprise rate (derived from the lexical frequency) and the contextual lexical surprise rate (derived from a deep neural transformers model) are of the same order of magnitude, but with the syllabic rate having the largest influence.

Among the seven factors investigated in this study, four pertain to information processing in the sense of Shannon’s theory of communication. Static and contextual lexical surprises are directly related to the participants’ linguistic expectations: both unusual words and sentence structures hinder the capacity to overcome the challenge caused by a high compression rate. Noteworthy is that phonemic and syllabic information rates also have an impact – albeit more limited – on comprehension, in addition to the lexical level. Previous studies highlighting the importance of information rate did not disentangle the syllabic rate from the syllable and lexical information. In the present study, we investigated the syllabic /phonemic functional loads, viz. the importance of correctly identifying the presented syllable /phoneme to access the target word. In other words, misperceiving a high functional load syllable /phoneme may lead to a wrong identification at the word level. Our study thus reveals the role of these phonemic and syllabic contrastive information once the lexical linguistic expectations are taken into account.

We also addressed whether in natural speech and at normal speed, the intrinsic statistics associated with each linguistic feature are already close to their channel capacity. Apart from contextual information, all other features’ generic statistics are below their respective channel capacity. Based on those results, we propose that contextual lexical surprise is an important constraint regarding the rate at which natural speech unfolds. Accordingly, speech production and perception can be envisioned as a dynamical information processing cycle, in which the speaker and the listener are two elements in interaction within one closed-loop converging system (Ahissar & Assa, 2016). While in this study we approach the question from the perception side, to delimitate the highest rate at which linguistic inputs can be processed, it would be of great interest to look at the same phenomenon from the production side and determine whether constraints imposed on speech comprehension have some equivalents in speech production. Related to this, investigating whether and which channel capacities can be extended by training could be a powerful way to optimise rehabilitation strategies in patients suffering from speech impairments.

Artificially compressing speech can lead to a degradation of the quality of the linguistic information. This can cause comprehension to drop as linguistic features may most efficiently be represented at their natural rates in the auditory system. However, previous work has repeatedly demonstrated that limitations in compressed speech comprehension are not due to limited capacities in acoustic information encoding. Neural activity recorded in the primary auditory cortex can indeed track the acoustic modulation rate even well outside of the intelligibility range (Nourski et al., 2009; Pefkou, Arnal, Fontolan, & Giraud, 2017). This feat is putatively rendered possible by the short temporal integration windows of early auditory areas (Giroud et al., 2020; Lerner et al., 2014; Poeppel, 2003). Conversely, the degraded comprehension of speeded speech is thought to arise from limitations of higher order brain areas in their speech-decoding capacities (Vagharchakian et al., 2012). A further argument in favor of this interpretation is that inserting delays between segments of highly compressed speech restores comprehension (Ghitza & Greenberg, 2009), highlighting the fact that is not a problem of stimulus encoding processing but rather a limitation in the time needed to decode the information present in the acoustic signal (Pefkou et al., 2017). By using time-compressed speech, we artificially increased the amount of information per time unit, leading to a drop in comprehension as a result of multilevel limited channel capacities, reflecting internal processes which can not keep up with the overflow of information. This saturation can be considered as analogous to attentional blink and psychological refractory period phenomena (Pashler, 1984; Raymond, Shapiro, & Arnell, 1992; Sigman & Dehaene, 2008) or more complex theoretical frameworks (S Marti, King, & Dehaene, 2015; Sébastien Marti & Dehaene, 2017), which suggests that the complexity of an integration operation defines its channel capacity. Our data are in accordance with this idea, as we showed that multilevel linguistic features predict accelerated speech comprehension performance. One question we can not answer is whether this is the result of a serial chain of processes or of competing parallel processes, or both. Further work using time-resolved measurements of comprehension could adjudicate between these concurrent hypotheses.

Finally, while we used meaningful sentences and words derived from large databases, due to experimental conditions, we artificially accelerated the spoken material to carefully control for speed variations. This controlled experimental task may seem somewhat unnatural but we show that the compressed speech gating paradigm is sensitive to linguistic features that have been shown to influence language processing in more classical experimental settings. Importantly this paradigm allows comparing in a generic framework different linguistic features from previously distinct subfields in the language domain. While the model approach comparison used in this work only affords relative conclusions, it undoubtedly paves the way for more thorough investigations of the effects of multilevel linguistic features on speech comprehension. Thanks to an innovative paradigm and stimuli selection procedure, our approach unifies a diverse literature under the unique concept of channel capacity. Our findings highlight the relevance of using both natural speech material (despite being more methodologically constraining) and a normative measurement framework to study speech comprehension. We hope that this work will settle the ground for further explorations of speech comprehension mechanisms at the interface of multiple linguistic research fields.

## Materials and Methods

### Participants

For experiment 1, 21 native French speakers (12 females, mean age 24.3 y, standard deviation ± 2.6, range [20, 30]) were recruited from Aix-Marseille University. For the second experiment, 21 French participants (11 females, mean age 22 y, standard deviation ± 1.6, range [20, 26]) were recruited online from Aix-Marseille University’s student group to perform the experiment through the FindingFive online platform. 20 French participants (12 females, mean age 25.5 y, standard deviation ± 5.7, range [20, 43]) took part in experiment 3. This experiment was also runned online thanks to the FindingFive platform. All participants reported normal audition and no history of neurological or psychiatric disorders. They provided informed consent prior to the experimental session. Participants received financial compensation for their participation. The experiments followed the local ethics guidelines from Aix-Marseille University.

### Stimuli

#### Speech stimuli

The stimuli in experiment 1 consisted of 251 monosyllabic French words drawn from a set of 1,100 monosyllabic words listed in the Lexique database (New et al., 2004). The stimuli in experiment 2 consisted in 100 seven-word-long French sentences drawn from a set of 14,000 seven-word sentences listed in the Web Inventory of Transcribed and Translated Talks database (WTI3, Cettolo et al., 2012). For both experiments, the text stimuli were then synthesized in auditory stimuli using Google Cloud Text-to-Speech (Google, Mountain View, CA, 2020, the female voice, “fr-FR-Wavenet-C”).

Using text-to-speech technology as opposed to naturally-produced speech has the critical advantage of controlling for the relevant linguistic features. Indeed, naturally produced speech displays variability across utterances in multiple linguistic characteristics (i.e., prosody, quality of phonetic pronunciation, phonemic duration, coarticulation, local speech rate, etc) (Miller, Grosjean, & Lomanto, 1984). On the contrary, synthetic speech remains highly consistent across utterances with the same sentence being always pronounced the same way. This point is highly important when assessing channel capacity, as the different words (Experiment 1) or sentences (Experiment 2) must be pronounced similarly to be able to estimate the impact of linguistic features on comprehension across stimuli.

Stimuli were selected on the basis of their characteristic linguistic features. For that, each stimulus at original speed was characterized by a vector composed of five features in experiment 1 and seven features in experiment 2. These linguistic features characterize the stimuli at different levels of processing, from acoustic to supra-lexical properties. Importantly, each feature was estimated in a number of units per second (i.e., in rate, or bit/s) to allow comparing their respective importance on speech comprehension (Coupé et al., 2019; Pellegrino et al., 2011; Reed & Durlach, 1998). The features were the following:

#### Acoustic modulation rate

it corresponds to the main acoustic modulation rate present in the speech signal. For each stimulus (words or sentences), the wideband envelope of the speech waveform was estimated (Chandrasekaran, Trubanova, Stillittano, Caplier, & Ghazanfar, 2009; Smith, Delgutte, & Oxenham, 2002) : the raw speech waveform was band-pass filtered into 32 frequency bands from 80 to 8,500 Hz with a logarithmic spacing, modelling the cochlear frequency decomposition. The absolute value of the Hilbert transform of each band-passed signal was extracted and summed across bands. The resulting envelope time-course was downsampled to 1000 Hz. Then, we used Welch’s method (Virtanen et al., 2020) to estimate the power spectral density of the envelope, resulting in a modulation spectrum between 1 and 215 Hz with a 0.1 Hz resolution. This was done for each stimulus. Finally, the center frequency of each spectrum was extracted by taking the global maximum value of each modulation spectrum. The acoustic modulation rate was expressed in Hz.

#### Phonemic rate

it corresponds to the number of phonemes presented per second. It was computed by dividing the number of phonemes (retrieved from the canonical pronunciation provided in the Lexique database (New, Pallier, Brysbaert, & Ferrand, 2004)) by the duration of the stimulus. The phonemic rate was expressed in Hz.

#### Syllabic rate

same as the phonemic rate but for syllables. It was also expressed in Hz.

#### Phonemic information rate

it measures how much information, defined by Shannon’s theory of communication, is carried by each phoneme (n=38). In order to approach this level from a perspective different from the lexical level described below, we adopted a methodology based on the contrastive role of the phonemes in keeping the words different in the French lexicon. For each distinct phoneme, its contrastive role was computed as its relative functional load (Oh, Coupé, Marsico, & Pellegrino, 2015). The functional load allows calculating the relative importance of a phoneme for a given language. More specifically, it quantifies its importance in terms of avoiding homophony keeping the words distinct in the lexicon, given their frequency of usage. The phonemic information rate is consequently defined for each stimulus as the sum of its phonemic functional loads divided by its duration. This feature was estimated from written data derived from the Lexique database. The phonemic information rate was expressed in bits per second.

#### Syllabic information rate

same as phonemic information rate but for syllables (n=3660). It was also expressed in bits per second.

#### Static lexical surprise rate

Derived from the lexical frequency, it measures the unexpectedness of a word without reference to the surrounding context. It was computed as the negative base 2 logarithm of the unconditional probability of a word -log2P(word), where P(word) is the lexical frequency of the word. The lexical frequency was the frequency of occurrence in the Lexique database. In experiment 1, the static lexical surprise was divided by the stimulus duration. In experiment 2, as stimuli were seven-word sentences, the static lexical surprise of each individual word composing the sentences was summed before dividing by the duration of the stimulus. The static lexical surprise was expressed in bits per second.

#### Contextual lexical surprise rate

Derived from a deep neural transformers model, it measures the unexpectedness of a word given the sentence context. It was computed as the negative base 2 logarithm of the conditional probability of a word -log_2_P(word|context), where P(word|context) is the probability of a word estimated by the french Bidirectional Encoder Representations from Transformers CamemBERT (Martin et al., 2020). This transformer network is a bidirectional- attention model that uses a series of multi-head attention operations to learn context-sensitive representations for each word in an input sentence in a self-supervised way by predicting a missing word given the surrounding contexts in large text corporas. We used the HuggingFace transformers Python package (Wolf et al., 2020) to access the pre-trained CamemBERT model with no further fine-tuning. Each individual sentence stimulus was passed through CamemBERT and the pooled output was averaged over the seven words contained in the sentence. This quantity was finally divided by the stimulus duration. As a context is needed to estimate the contextual lexical surprise, it was only computed for experiment 2, where stimuli are sentences. The contextual lexical surprise was expressed in bits per second.

### Procedure and Paradigm

#### Orthogonalisation procedure to select the stimulus sets

In order to avoid collinearity issues due to correlations between features across stimuli, we developed a custom-made leave-one out iterative algorithm to select stimuli with low correlation between features. The algorithm starts with the complete original database (1,100 words in experiment 1 and 14,000 sentences in experiment 2) and computes the correlation between each pair of features (5-7 features, 10-21 correlations in total in experiment 1 and 2 respectively). Then, the algorithm performs a leave-one-out procedure: it removes one stimulus, recomputes the correlation matrix on this reduced set and estimates the specific contribution of the one stimulus on the original correlation matrix, by comparing the correlation matrices of the full and reduced stimuli sets. This processing step is repeated until all items have been removed once. The 10 percent stimuli that led to the most significant increase in correlation across features are discarded. The algorithm then iterates on this newly selected reduced stimuli set. The algorithm stops when the number of stimuli is equal to 251 (words) in experiment 1 and 100 (sentences) in experiment 2. A last check ensured that the correlations between features were all below 0.15.

#### Representativeness of the selected stimulus sets

The representativeness of the final selected stimulus sets in comparison to the original datasets was assessed for each feature. This was performed to ensure that any theoretical conclusions derived from the results obtained from a limited subset of stimuli could generalize to a larger corpus-based dataset. To do so, we computed the value of the features for the complete datasets, hence providing a relatively good estimate of the ecological distribution of each feature. Two indexes were computed to control that each feature’s distribution in the selected stimulus sets was similar to its distribution of the original datasets: i) the ratio between the means, ii) the ratio between the variances. A value close to one for both indexes indicates a good match between the distributions in the original dataset and in the selected stimulus sets. Finally, the correlation matrices between the features in the selected stimulus sets and the features in the original datasets were compared.

#### Time compression

Time compressed versions of each stimulus were created. The audio waveforms were linearly compressed at rates 1, 2, 2.2, 2.5, 2.9, 3.5, 4.3, 5.6, 8 and 10 of the original recording in experiment 1, at rates 2, 2.5, 3, 3.5, 4, 4.5 and 5 in experiment 2 and finally at rate of 3.5 for experiment 3. A compression rate of 2 indicates that the duration of the time-compressed version of the audio file is equal to half of the natural duration. The compression rates in experiment 2 were adjusted on the basis of the results of experiment 1. The PSOLA algorithm implemented in the Parselmouth Python package based on PRAAT (Boersma, 2001; Jadoul, Thompson, & de Boer, 2018; Moulines & Charpentier, 1990) was used to modify the duration of the audio stimulus without altering the original pitch contour. Audio stimuli were normalized in amplitude and digitized at 44.1 KHz. This resulted in 2510 audio stimuli (251 words x 10 compression rates) in experiment 1, 700 audio stimuli (100 sentences x 7 compression rates) in experiment 2 and 100 audio stimuli (100 sentences x 1 compression rate) in experiment 3. A manual check was performed to ensure that the compression procedure did not insert salient quirks.

One necessary prerequisite of our experiment is that across presentation rates all the investigated acoustic and linguistic factors are uniformly modified (i.e., that time-compression does not impact a particular feature more than the others). Previous experimental work has shown that artificially time- compressed speech and natural fast speech are qualitatively different. Indeed, in the first case, the spectral content is exactly similar but the duration of the utterance is reduced. This results in an uniform modification of all spectral and temporal details. In the second case, due to restrictions on articulation, the signal is affected non-uniformly (Guiraud et al., 2018; Janse, 2004). In addition, the idea of using the modified gating paradigm was to present to the participants at each compression rate exactly the same overall quantity of information, albeit delivered at different speed/rate, so that the channel capacity of each factor can be estimated. Hence it was crucial that the material was exactly similar across compression rates, except for the time dimension.

#### Paradigm

All three behavioral experiments consisted in a modified version of the gating paradigm (Grosjean, 1980) using time-compressed speech stimuli.

In experiment 1, participants were presented with 10 time-compressed versions of isolated words. Each trial consisted in the successive presentation of different time compressed versions of the same audio stimulus, in an incremental fashion, starting with the most compressed version of the stimulus (gate n°1) and ending with the least compressed version (either gate n°10). After each audio presentation, participants were asked to type on the keyboard what they heard and then to press enter to continue to the next gate.

Experiment 2, was similar to experiment 1, apart from the fact that participants were presented with 7 time-compressed versions of seven-word sentences. Each trial thus ends at gate n°7, following the presentation of the least compressed version of the sentence. In experiment 2, participants were required to repeat in the microphone at each gate what they heard and then to press enter to continue to the next gate.

Experiment 3 was similar to experiment two except that only one time compressed version (x 3.5) of each sentence was presented per trial.

In all experiments, participants were instructed that each auditory stimulus was meaningful and difficult to understand at the highest compression rates. In order to get familiarized with the task, participants completed three practice trials before the experiment. Experiments 1 and 2 were composed of two sessions of approximately 50 minutes each. The sessions included several breaks for the participants to stay vigilant and focussed throughout the experiment. Each participant was presented with the stimuli in a pseudo-randomized order. The experiments were self-paced and there were no time constraints. The two sessions were performed at most one week apart. Experiment 3 took 25 minutes to complete. The paradigm used in all experiments incorporated a transcription task which required participants to explicitly recognise, recall, and either reproduce each isolated word or each word of the sentence. It provided a fine-grained accuracy measure associated with focused and extensive linguistic processing. A pilot study was performed to properly select the multiple compression rates in the first experiment. For the second experiment we adjusted the compression rate based on the first experiment and another pilot study. Overall, the range of values of the different compression rates have been appropriately chosen and capture the sigmoid shape of our psychometric data.

#### Experimental setup

Experiment 1 was implemented in Python with the expyriment package (Krause & Lindemann, 2014) and run on a ASUS UX31 laptop. The program presented the audio stimuli binaurally at a comfortable hearing level via headphones (Sennheiser HD 250 linear) and recorded the participants’ written responses. Participants came to the laboratory and performed the two sessions in an anechoic room. Due to the Covid-19 outbreak, two different sets of participants undertook experiment 2 and 3 online via the experimental platform FindingFive (FindingFive, 2019). The procedures were the same except that participants were instructed to record their answers with a microphone (instead of typing them) to optimize the duration of the experiment.

### Data analyses

#### Data scoring

Speech comprehension was scored 1 if the response was correct (grammatical errors were allowed) and 0 if the response was incorrect or if no answer was given. In experiment 2 and 3, participants’ audio responses were first transcribed using Google Cloud Speech-to-Text (Google, Mountain View, CA, 2018) and checked manually for mistakes or inconsistencies.

#### General linear mixed model (GLMM) analysis

Participant’s responses (0: incorrect, 1: correct) were analyzed using Generalized Linear Mixed Models (GLMM; (Quené & van den Bergh, 2008) with a logistic link function using the lme4 package (Bates, Mächler, Bolker, & Walker, 2015) in R (version 3.5.1,Team, n.d.). The datasets were composed of 52,710 responses in experiment 1 (21 participants x 251 words x 10 compression rates) and 102,900 responses in experiment 2 (21 participants x 100 sentences x 7 words x 7 compression rates). Acoustic modulation rate, phonemic rate, syllabic rate, phonemic information rate and static lexical surprise were entered as fixed effects in experiment 1. Participants and compression rates were entered as random effects. The model was expressed as follows in lme4 syntax:

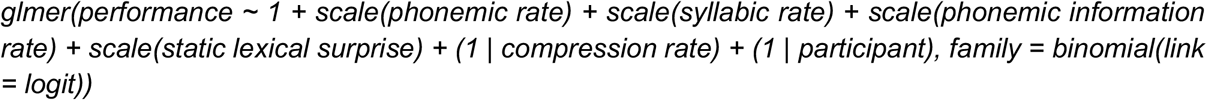

In experiment 2, the model was the same except that syllabic information rate and contextual lexical surprise were added as fixed effects. The model was:

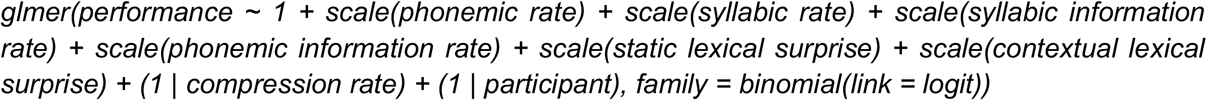

No interaction terms were estimated in the models. First, models including all the possible interactions failed to converge. Second, converging models that included a subset of interactions only very marginally increased the percentage of variance explained in the behavioral responses (marginal and conditional R^2^). These latter are well and best captured by the main effects.

Post-hoc comparisons between the resulting estimates associated with each feature were conducted using the glht function from the multcomp package in R (Hothorn, Bretz, Westfall, & Heiberger, 2016). All p-values reported were corrected for multiple comparisons using the Holm correction.

#### Comprehension point determination

For each stimulus, the comprehension point was estimated. It is defined as the compression rate at which participants reached a 75% correct response performance, as predicted by a logistic function. Fitting procedures were performed in R using the glm function from lme4 package (Bates et al., 2015).

#### Linear mixed model (LMM) analysis

Comprehension points were analyzed using linear mixed models (LMM). This complementary statistical analysis aimed at characterizing the relationship between the values of each feature at normal speed and the comprehension points. The rationale was that if they impact comprehension, the feature values at normal speed are predictors of the compression rate at which comprehension shifts from incorrect to correct. Whereas, in the GLMM analysis, all behavioral responses were entered in the model, the current analysis exploits only the comprehension point in each trial. The final datasets were composed of 5,271 comprehension points in experiment 1 (21 participants x 251 words) and 2,100 comprehension points in experiment 2 (21 participants x 100 sentences). Acoustic rate, phonemic rate, syllabic rate, phonemic information rate and static lexical surprise were entered as fixed effects in experiment 1. Participants and compression rates were entered as random effects. The model was:

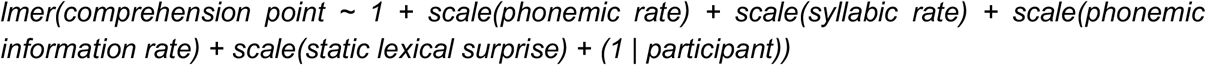

In experiment 2, the model was the same except that syllabic information rate and contextual lexical surprise were added as fixed effects. The model was:

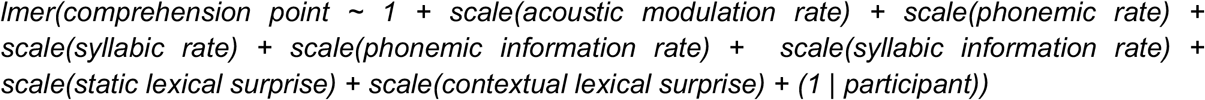

#### comparison of regressors across experiments 1 and 2

Following the method recommended by (Paternoster, Brame, Mazerolle, & Piquero, 1998), we statistically assessed the significance of the difference between the multiple regressors across experiments 1 and 2 in an unbiased way using their standardized estimates and standard error to the mean. Moreover, after having transformed the resulting Z-scores (standard normal distribution) into p-values, we additionally applied a Holm-correction for multiple comparisons. From the resulting statistics, we assessed, for each linguistic feature, potential significant differences between experiments 1 and 2.

#### Determination of channel capacity associated with each linguistic feature

The processing of each linguistic feature was modeled as a transfer of information through a dedicated channel. Channel capacity is defined as the maximum rate at which information can be transmitted. For each feature, it was estimated using the comprehension point and defined as the value of the feature at the comprehension point.

#### Overlap between channel capacity and generic features distributions

The overlapping R- package (Pastore, 2018) was used to compute the percentage of overlap between the values of the channel capacity associated with each feature and their generic distribution in the stimulus set at normal speed. The method divides the density distribution into intervals and computes the cumulative sum of minimum values per interval. The result can vary between 0 and 1, where 1 indicates that the two distributions are identical and 0 indicates a complete absence of overlap. The percentage of overlap between feature distributions reveal which feature is already near the upper limit of speech comprehension at normal speed, potentially limiting our ability to cope with higher speed speech.

#### Model validation

All models were fitted in R (version 3.5.1, (R core, 2020)) and implemented in RStudio (Racine, 2012) using the lme4 package (Bates et al., 2015). Fixed effects were z- transformed to obtain comparable estimates (Schielzeth, 2010). Visual inspection of residual plots was systematically performed to assess deviations from normality or homoscedasticity. Variance inflation factors (VIF) were also checked to ensure that collinearity between fixed effects was absent. Overall, VIF values were generally close to one and no deviations from model assumptions were detected. We tested the significance of the respective full models as compared to the null models by using a likelihood ratio test (R function anova). Goodness of fit of the models were evaluated and reported using both the marginal and conditional R^2^.

#### Data availability

Numerical data supporting this study will be available on GitHub: https://github.com/DCP-INS/

#### Code availability

Codes to reproduce the results and figures of this manuscript will be available on GitHub: https://github.com/DCP-INS/

## Acknowledgments

We thank all participants; Johanna Nicolle, François-Xavier Alario and all the colleagues from the DCP team at the Institut de Neurosciences des Systèmes for useful discussions; Yannick Jadoul for help with the Parselmouth python package and Ting Qian from FindingFive for extensive assistance and advice.

## Supplementary Figures

**Figure Supplementary 1.**
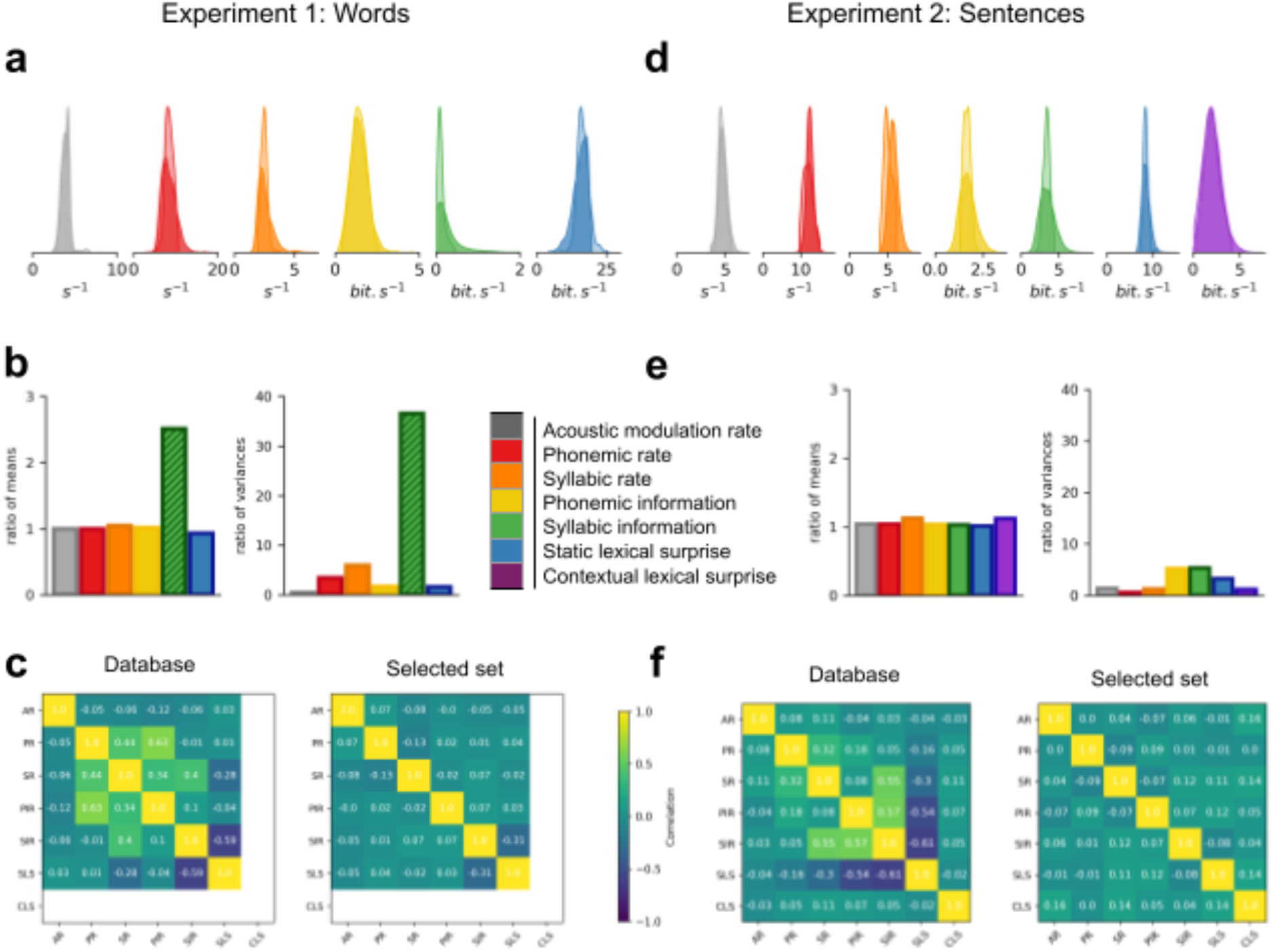
Description of the linguistic features in the original database and selected stimulus set, for experiments 1 (a-c) and 2 (d-f). **a**,**d)** Distribution of the linguistic features in the original database (dark colors) and selected stimulus set (light colors), at original speed. **b**,**e)** Ratios of means (left) and variance (right) across stimuli, between the selected stimulus set and the database. **b)** Striped (green) bars highlight an outlier linguistic feature in experiment 1, for which the selected stimulus set is not representative of the original database. **c**,**f)** Correlation matrices between linguistic features in (left) the original database and (right) selected stimulus set. The selection procedure ensured that low correlations (all r < 0.15) across stimuli were present between features in the selected stimulus sets (see Methods). AMR: acoustic modulation rate, PR: phonemic rate, SR: syllabic rate, PIR: phonemic information rate, SIR: syllabic information rate, SLS: static lexical surprise and CLS: contextual lexical surprise.

**Figure Supplementary 2.**
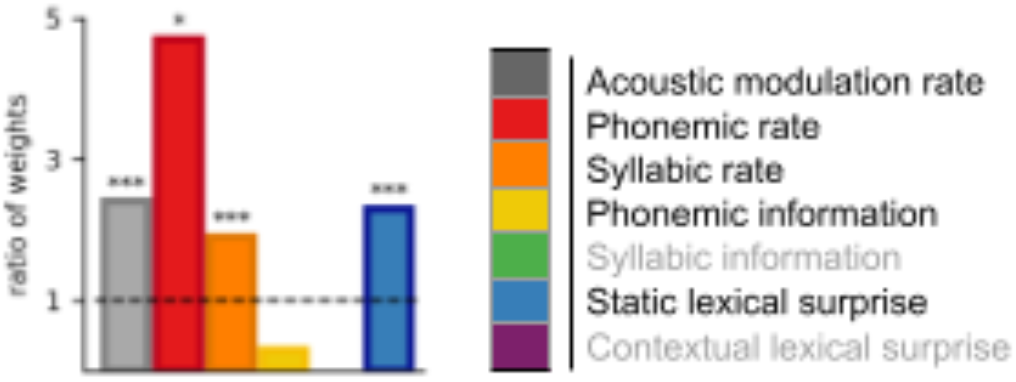
Comparison of experiments 1 and 2. Ratios of the standardised weights estimated from experiments 1 and 2. P-values are estimated after Paternoster et al. (1998). *p < 0.05; ***p < 0.001.

**Figure Supplementary 3.**
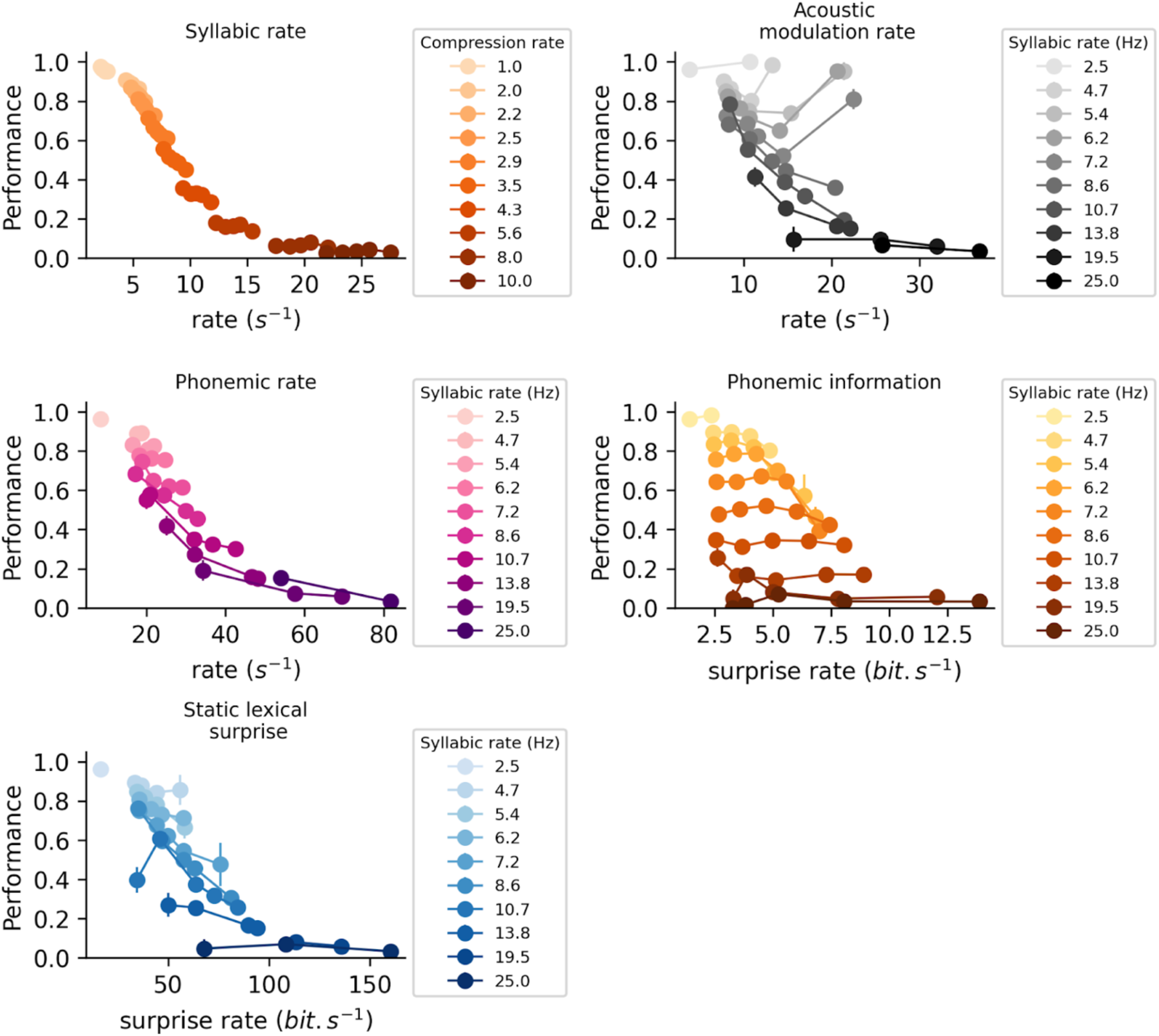
Experiment 1. Comprehension performance as a function of the different linguistic features. Performance is expressed in proportion of correct responses. **Upper left panel:** Performance sorted as a function of the compression rate (colorscale) and the syllabic rate (y-axis). **Other panels:** Performance sorted as a function of the syllabic rate (colorscale) and the different linguistic features (y-axes). Data were sorted as a function of the syllabic rate as this feature had the strongest impact on comprehension performance (see Fig. 3) and could thus hide the impact of the other features in this visualisation.

**Figure Supplementary 4.**
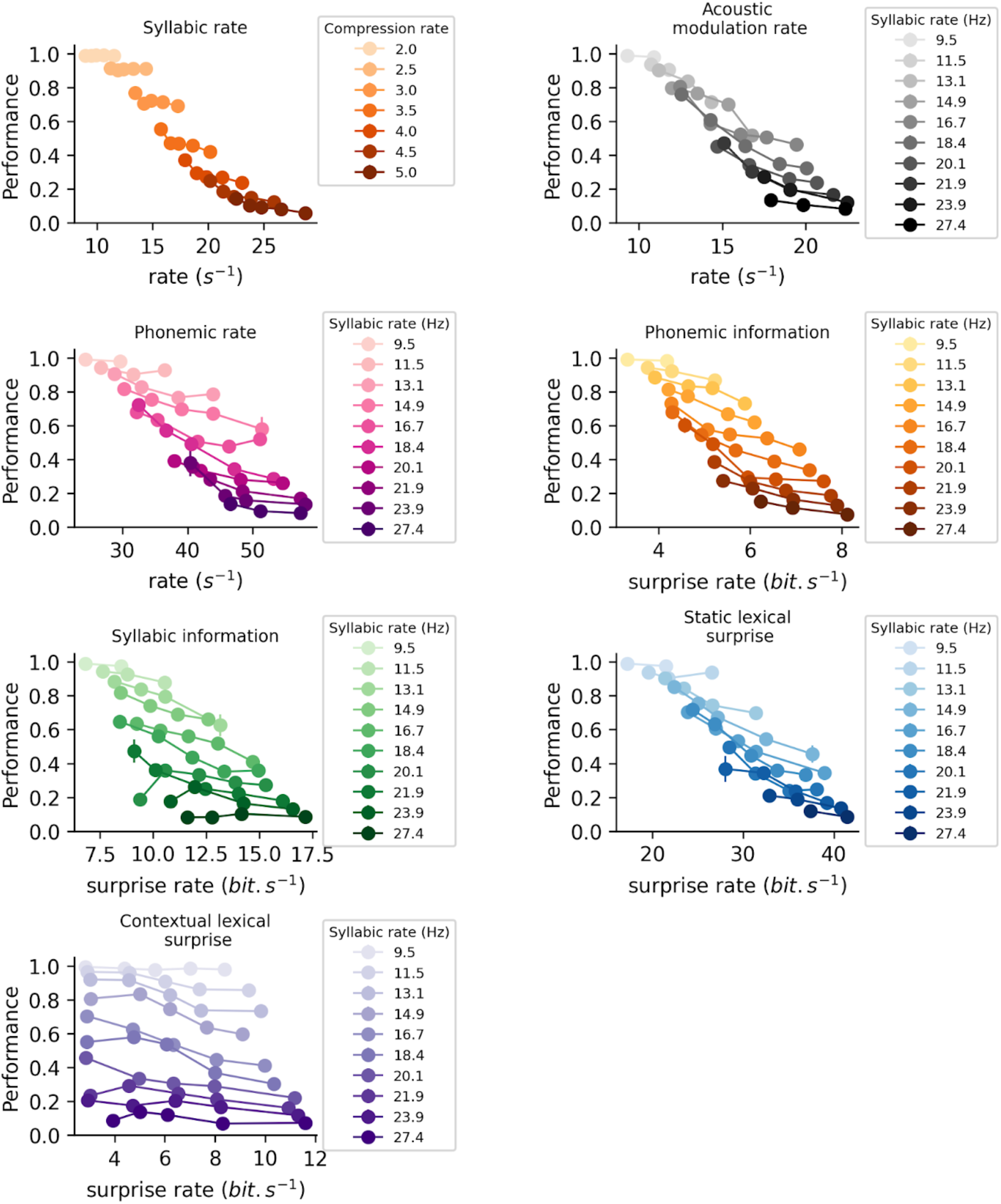
Experiment 2. Comprehension performance as a function of the different linguistic features. Performance is expressed in proportion of correct responses. **Upper left panel:** Performance sorted as a function of the compression rate (colorscale) and the syllabic rate (y-axis). **Other panels:** Performance sorted as a function of the syllabic rate (colorscale) and the different linguistic features (y-axes). Data were sorted as a function of the syllabic rate as this feature had the strongest impact on comprehension performance (see Fig. 3) and could thus hide the impact of the other features in this visualisation.

**Figure Supplementary 5.**
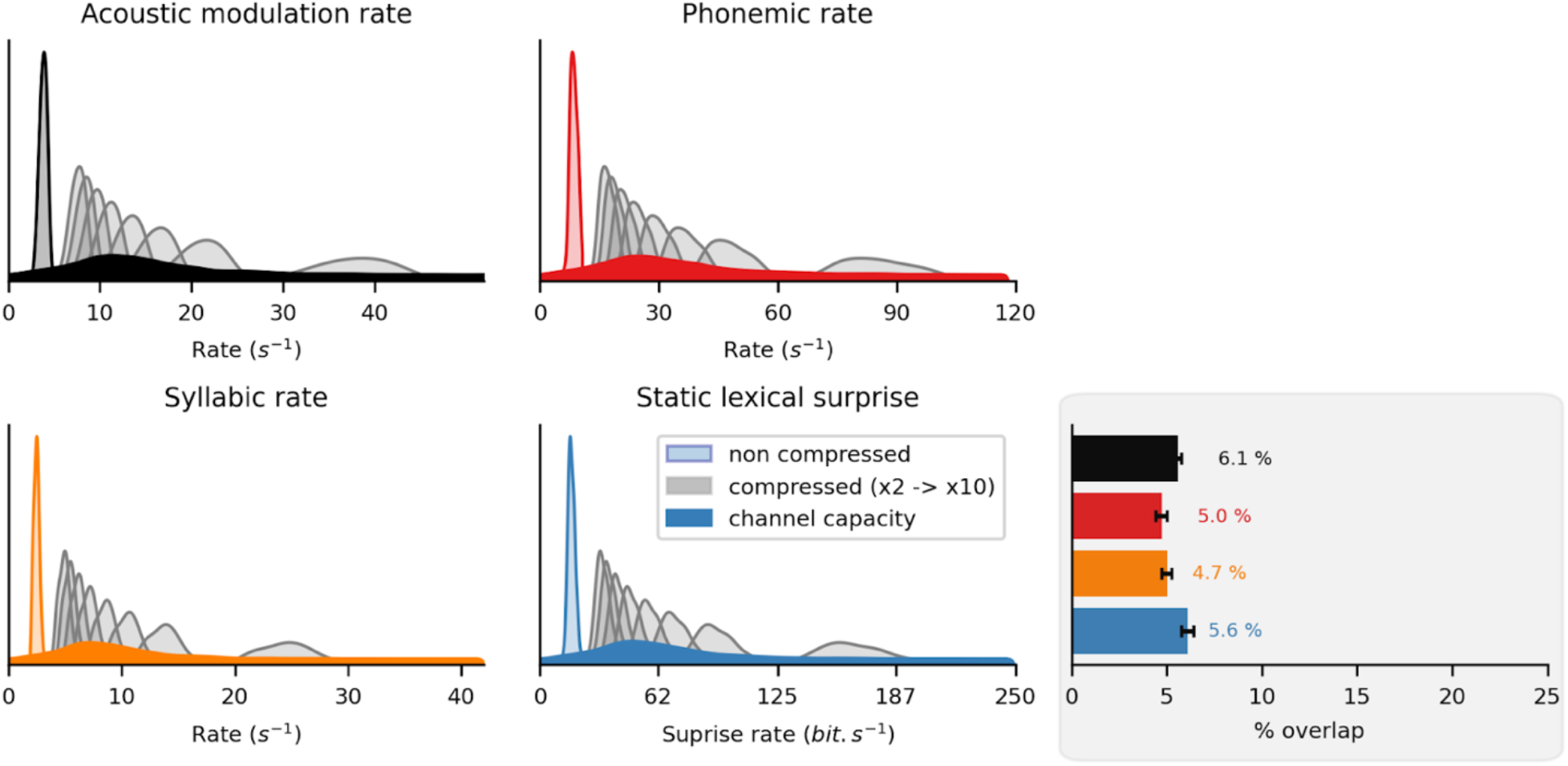
Experiment 1 (words). Overlap between the linguistic channel capacities and their generic distribution in the stimulus set. Distribution of the linguistic features in the selected stimulus set at original speed (non-compressed, lighter color) and at the different compression rates (in grey). Superimposed is the corresponding estimated channel capacity (see Fig. 5; darker color). **Lower right (grey panel):** Overlap ratio between the channel capacity associated to each linguistic feature and its distribution at original speed. Error bars indicate standard error of the mean across participants.

